# The kin-selected context of duelling in horned aphids: cooperation or conflict?

**DOI:** 10.1101/2025.01.29.635201

**Authors:** Keigo Uematsu, Man-Miao Yang, William Foster

## Abstract

In one-on-one (dyadic) contests in group-living animals, relatedness is of potentially critical importance in determining the relative value of the resources for which the individuals are fighting. We investigated the function of dyadic contests over access to a food resource (plant phloem) in the horned aphid *Astegopteryx bambusae*, by observing competing aphids in the field and genotyping them with microsatellite markers. Each aphid colony consists of a small number of clones, creating levels of relatedness that can be both high and variable. The contests involve only low costs in terms of time or injury.

Our results indicate that there is no kin-discrimination in the duelling pairs of aphids. Genetic relatedness between duelling pairs did not differ significantly from that of randomly selected pairs and showed no association with contest duration or outcome. The mean relatedness between a duelling pair was 0.79 ± 0.12 (mean ± SD, *N* = 75), with 56% (42/75) of duels occurring between clonal pairs.

We suggest that the duels in these *Astegopteryx* aphids are not an aggressive fight for resources between different genotypes, but rather a low-cost method by which the aphids assess each other’s reproductive value. 83% (50/60) of the contests between aphids of different ages were won by the aphid that was older, and therefore of greater reproductive value, providing an indirect fitness benefit for the losing younger individual. Younger nymphs, when attacked by older aphids, yield feeding sites altruistically and gain inclusive fitness benefits: this provides a compelling selective context for the evolution of the young, rather than old, altruistic soldiers that are observed in the open colonies of many cerataphidine species.

## Introduction

Animal combat has always fascinated biologists and the adaptive interpretation of one-to-one (dyadic) fighting has played a pivotal role in clarifying our understanding of the fundamental concepts of evolutionary adaptation: the level at which selection acts and the development of game-theoretic models of animal behaviour (Hardy & Briffa, 2013; Parker, 1974). Dyadic fights have been studied extensively using theoretical models and empirical tests that attempt to determine how the two contestants might assess each other’s combat ability, the relative value of the indivisible resource they are fighting for, and the likely costs of the fight (Green et al., 2021; Kokko, 2013).

In group-living animals, however, an additional factor - genetic relatedness - can critically affect the selective landscape of a dyadic fight (Croft et al., 2021; Hardy & Mesterton-Gibbons, 2023; Taborsky et al., 2021). If you are fighting a relative, kin selection will mean that your actions will affect not only your fitness but also that of your opponent. For example, reproductive skew in a group, in which dispersal is limited, promotes kin selection for harmful behaviour by dominant individuals and helping behaviour by subordinates (Johnstone, 2008). In addition, when there is significant heterogeneity in age or fecundity, differences in reproductive value might encourage an individual of low reproductive value to help one of higher reproductive value (Hasegawa & Kutsukake, 2019; Rodrigues & Gardner, 2022; Uematsu et al., 2013). Intense, apparently antagonistic, contest behaviour can promote cooperation by enabling the swift transfer of an essential, indivisible resource, thereby increasing the inclusive fitness of the helper. To clarify the function of dyadic contests in a kin-structured group, it is therefore vital to understand the genetic relatedness of the contestants in their natural habitat.

Animals that live in groups and reproduce clonally provide an ideal context in which to spotlight the importance of genetic relatedness on dyadic fighting, since the variation in levels of relatedness is about as high as we can expect to see in nature: ranging from those that are undoubtedly from the same clone to those that are entirely unrelated. Clonality, which occurs in about two thirds of the metazoan phyla (Hughes, 1989), gives rise to groups of two main sorts: modular colonies, where the clonal individuals are functionally integrated, as in many colonial benthic invertebrates such as Anthozoa and Bryozoa; and unitary groups, made up of independent clonal individuals, such as aphids and cladocerans (Bell, 1982). Fighting behaviour has been studied in a number of modular benthic invertebrates, although the fights are often between several individuals in two opposing colonies rather than strictly dyadic fights. Fighting between independent clonal individuals has been relatively little studied, although such fights should provide unusually clear situations in which to assess the importance of relatedness to the outcome of dyadic contests.

We report here on a species of aphid, *Astegopteryx bambusae* (formerly *A. bambucifoliae* (Li et al., 2019); tribe Cerataphidini), which provides an exceptionally amenable system in which to look at the kin-selected context of dyadic fighting. Several cerataphidine species exhibit dyadic fighting and it is currently unclear why it is only in the Cerataphidini, containing just 116 of more than 5000 species of Aphidoidea (Favret, C., 2024. Aphid Species File. Retrieved on 3 December 2024. Available from URL: http://Aphid.SpeciesFile.org), that fighting for a feeding site has evolved. The answer almost certainly lies in the choice of secondary host-plant (always monocotyledons) in these species and the fact that these aphids tend to feed on relatively mature parts of their host-plants. We know from previous work on this aphid and a closely related species *A. minuta* what the aphids are fighting for - access to a feeding site on the leaves of bamboo (Aoki & Kurosu, 1985; Foster, 1996). This prize is worth the fight. Aphids and plants have been in an arms race over access to phloem for thousands of years and, as a result, it is always very time- consuming for aphids to find and insert their stylets into the vascular bundle. Morris and Foster (Morris & Foster, 2008), however, showed – using EPG (electrical penetration graphs) - that an aphid that gained access to a ready-made feeding site by winning a fight was able to access the phloem within a matter of minutes. Aphids that did not fight but inserted into an abandoned feeding site took about 20 minutes longer to find the phloem. However, this is likely a significant underestimation of the potential risks and costs of avoiding conflict. Although we lack a precise measure of how long it would take one of these aphids to locate and establish its own feeding site, data from other non-fighting species suggest it could take several hours.

This aphid system provides a number of other important practical advantages. The duels are remarkably stereotypical and have already been well described (Aoki & Kurosu, 1985; Foster, 1996): the attacker is always a free-walking aphid which attacks a feeding aphid, which turns to face the attacker, without withdrawing its stylets, and raises its abdomen and lowers its head to shield itself against the attacker (Fig. 1; Supplementary Video S1). As the contest escalates, the attacking aphid clasps the opponent’s body with its forelegs, while the defender then raises its abdomen forward to lean on the attacker, performing a headstand. The costs of the fight seem small: they last for only about 3 minutes, and neither the horns nor the legs have ever been observed to damage the defender. The aphid colonies of this species are known to contain a variable number of clones (from 1- 20) (Uematsu et al., 2023): the aphids are therefore able to interact both with aphids from their own and from alien clones. The fights are easy to observe both in the field and laboratory; they occur frequently under natural conditions. The aphids can be allotted to one of five age classes (1^st^ – 4^th^ instar, adult), which is a decent proxy for size and a measure of resource holding potential.

**Figure 1.**
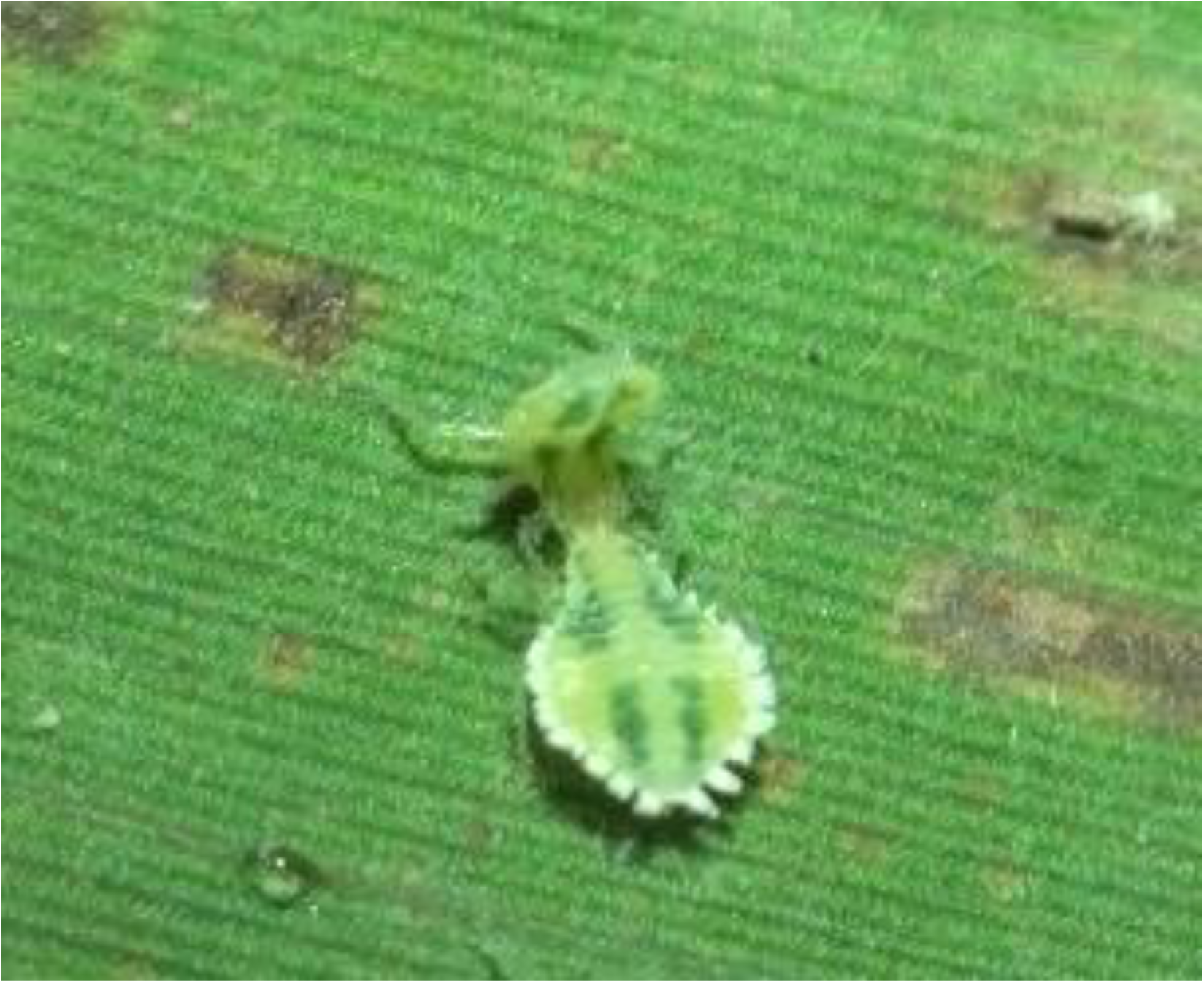
Head-butting in *A. bambusae* aphids. An attacker (front) butts the head of the defender (back) and clasps its body. The defender raises its abdomen and performs headstand. See also Supplementary Video S1.

Without knowing the pairwise relatedness of the fighting aphids it is not possible to assess the functional significance of what is clearly a common, routine behaviour in this and some other aphid species. A previous study, using newly developed microsatellite markers, showed that genetic relatedness within colonies of *A. bambusae* was high (0.52 on average) (Uematsu et al., 2023), indicating that aphids will frequently interact with members of the same clone. It is entirely possible that the duels are not an aggressive fight but a means of mutually communicating the relative reproductive value of the two aphids: younger aphids might initiate feeding sites and subsequently yield them to older aphids (Carlin et al., 1994; Foster, 1996).

This study aims to answer the following questions.

1. Does an aphid’s size influence (a) whether or not it behaves as an attacker and (b) the size of aphid that it attacks?
2. Does genetic relatedness affect which aphids are targeted by an attacker?
3. Is the duration and outcome of contests influenced by the relative size and/or the genetic relatedness of the contestants?
4. What is the function of the fighting behaviour?

## Materials and methods

### Sampling in the field

We defined a “colony” of *A. bambusae* as a group of aphids located on the same leaf of the host plant *Dendrocalamus latiflorus*. We observed and sampled *A. bambusae* colonies on 20, 21 and 29 December 2013 and 23 and 24 April 2014 at Sun Moon Lake, Nantou County, Taiwan. Each colony of *A. bambusae* was observed for a duration of 30 minutes, focusing on the occurrence of the butting behaviour. When an individual aphid (attacker) displayed butting behaviour towards another aphid (defender), the age categories of both the attacker and the defender (first-instar, 2nd instar, 3rd instar, 4th instar, adult) based on their body size, and the duration of the competition were recorded. We defined the defender’s victory when the attacker ceased butting and left, and the attacker’s victory when the defender withdrew its mouthparts and vacated the feeding site. The time taken for the competition was recorded in minutes. The competing attacker and defender were separately collected by the observer (KU) and subsequently preserved in 99% ethanol by an assistant. After 30 minutes, the entire colony was preserved in 95% ethanol. In total, we observed 111 competing pairs from 23 bamboo colonies (16 colonies in December 2013 and 7 colonies in April 2014) and collected 75 pairs from 19 colonies for genotyping. In the laboratory, we determined the age category of the competing aphids and counted the total number of aphids in each colony, as well as the age category of whole colony members of 11 colonies under a dissecting microscope. In addition to the competing aphids, 5 – 12 individuals from each of 22 colonies were selected for DNA extraction and genotyping. In total, 370 individuals of *A. bambusae* were genotyped.

### DNA extraction

DNA extraction was carried out using a glass milk extraction method in a microtitre plate as described in Amos et al. (Amos et al., 2016). Each aphid was crushed with a plastic pestle or a pipette tip in 40 μl of lysis solution (10 mM Tris HCl (pH 8.0), 1 mM EDTA, 1% SDS) with proteinase K and incubated at 56°C for 12 hours. The DNA was adsorbed onto flint glass particles in the presence of a 3× excess of 6 M NaI. Following two ethanol washes, the DNA was eluted in 50 µl of low TE buffer.

#### Genotyping

Polymerase Chain Reactions (PCRs) were conducted using Type-it Microsatellite PCR Kit (Qiagen) in 10 µl reactions. PCR conditions were as follows: an initial denaturing step of 5 min at 95°C, followed by 35 cycles of 95°C denaturation for 30 sec, 60°C annealing for 90 sec, 72°C extension for 60 sec, followed by a final extension step for 30 min at 60°C. Six microsatellite primers developed in the previous study (Uematsu et al., 2023)were used to generate multilocus genotypes. One primer from each pair was labelled with a fluorescent dye (6-FAM / HEX / NED) as described in Table 1. Following amplification, the PCR product was mixed with the GeneScan 500 LIZ Size Standard (Applied Biosystems) and subjected to fragment analysis. Allele calling was performed using GeneMapper software (Applied Biosystems) and manually checked. Samples with ambiguous genotype were re-run twice and the consensus genotype was taken forward for analysis.

**Table 1.**
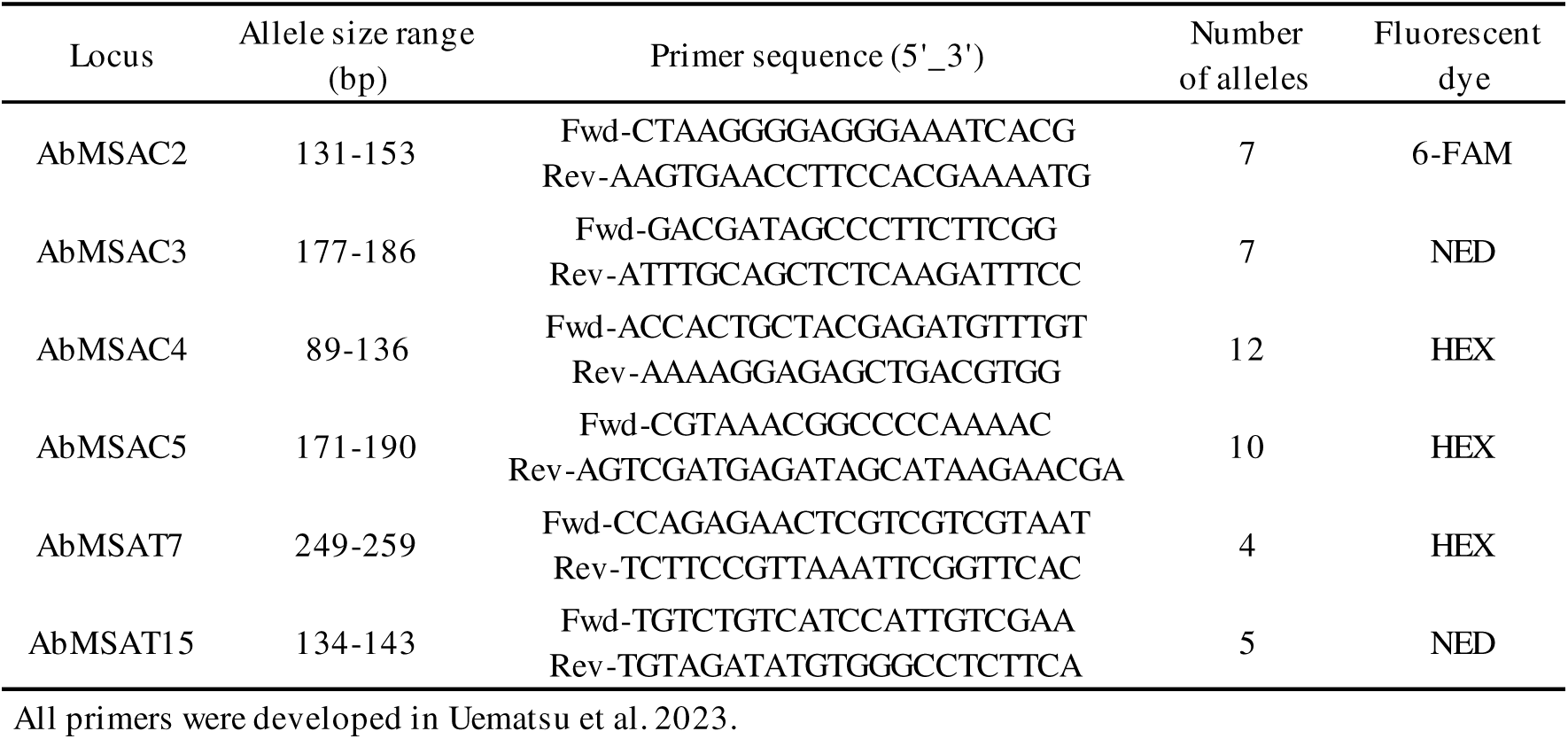
Polymorphic microsatellite loci with the primer sequences in *A. bambusae*.

#### Data analysis

All statistical analyses were conducted using the R software (R Core Team & Team, 2022). To investigate that the age distribution of both the attacking and defending aphids is not randomly chosen from the same colony, we generated 10,000 bootstrapped sets from the same colony with replacement. Under the null hypothesis, the observed age distribution should follow the distribution from resampled sets; under the alternative hypothesis (i.e. age-biased attack or defence), the observed age distribution should be significantly different.

To investigate the determinants of butting duration, separate generalized linear mixed models (GLMMs) were employed using the ‘glmer’ command from the lme4 package (Bates et al., 2015). Explanatory variables included age difference, age of attacker, age of defender, and the contest outcome (attacker’s win or defender’s win). The model selection was based on the lowest AIC, with the variance inflation factor (VIF) ensuring no multicollinearity between the explanatory variables (Zuur et al., 2010). Colony identity was set as a random effect. The significance of the fixed effects was determined based on Wald *Z* statistics and *P* values provided by ‘glmer’.

Different clones were identified using their multilocus genotypes. Pairwise relatedness values among all individuals were calculated following (Queller & Goodnight, 1989) using Kingroup (Konovalov et al., 2004) software. Average relatedness values between competing pairs and randomly selected pairs from the same colony were compared by generating 10,000 bootstrapped sets with replacement, assessing the significance of mean relatedness differences.

## Results

### Observation of butting behaviour in the field

We observed and analysed head-butting in twenty-three colonies of *A. bambusae*. The details of the behaviour were as described in previous observations on *Astegopteryx* species (Aoki & Kurosu, 1985; Foster, 1996). The total number of aphids in the colonies ranged from 82 to 794 (mean ± SE = 377 ± 36). The number of duelling pairs of aphids observed during the 30- minute period ranged from 0 to 14, showing no significant correlation with colony size (*t*_21_ = 0.77, *P* = 0.45). The average percentage of aphids in a colony that were observed to be attacking was 3.1 ± 0.6 (mean ± SE, *N* = 75) per hour.

### Relationship between aphid age and fighting behaviour

Of the 111 aphids observed to be attacking other aphids in 23 colonies, only 16 (14%) were 4^th^ instars or adults, the remaining 95 (86%) were 1^st^ – 3^rd^ instars (Supplementary Figure 1). This suggests that the younger aphids are more important than the older ones in initiating contests, but it could simply just be that they are more abundant in the colonies. We therefore measured the numbers of aphids of different ages within 11 colonies and compared this with observed numbers of aphids that behaved as attackers in these same colonies (Fig. 2). The distribution of attacking behaviour is clearly not random, with the second and third instars being more likely, and the 1^st^ instars and adults less likely, to behave in attack mode. Younger instars were more likely to lose when they were defenders (Cochran-Armitage trend test, χ^2^_1_ = 19.2, *P* < 0.0001; Table A1).

**Figure 2.**
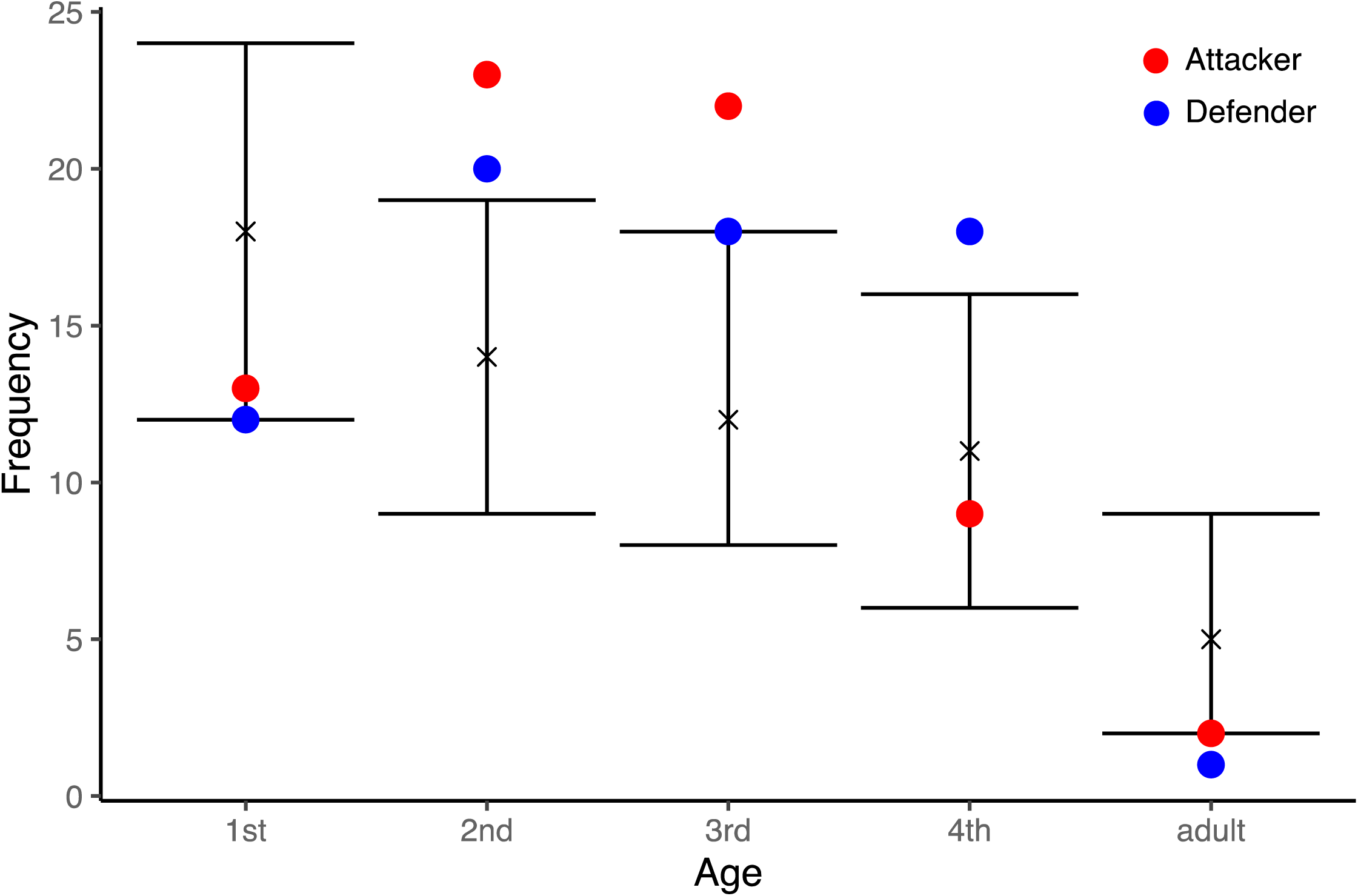
The frequency of butting pairs. Plots show the means and 95% bootstrap confidence intervals from 10,000 bootstrapped sets of the frequency distribution of the five age categories (1st to 4th instar and adult) within the observed colonies. Filled circles show the observed frequencies of attackers (red) and defenders (blue).

### Genetic relatedness between competing aphids

Average pairwise relatedness within a colony ranged from 0.06 to 1.0 (mean ± SD = 0.76 ± 0.08, *N* = 22, Fig. 3a), showing no significant correlation with colony size or butting frequency (vs. colony size: *t*_20_ = 1.68, *P* = 0.11; vs. frequency: *t*_20_ = 0.54, *P* = 0.60). The pairwise relatedness between the competing pairs (mean ± SD = 0.79 ± 0.12, *N* = 75) in the 19 sampled colonies was not significantly different from the distribution of relatedness values between randomly chosen pairs from across the same 19 colonies, weighted according to the number of fighting pairs sampled from each of these colonies (two-tailed bootstrap test, *P* = 0.59; Fig. 3b).

**Figure 3.**
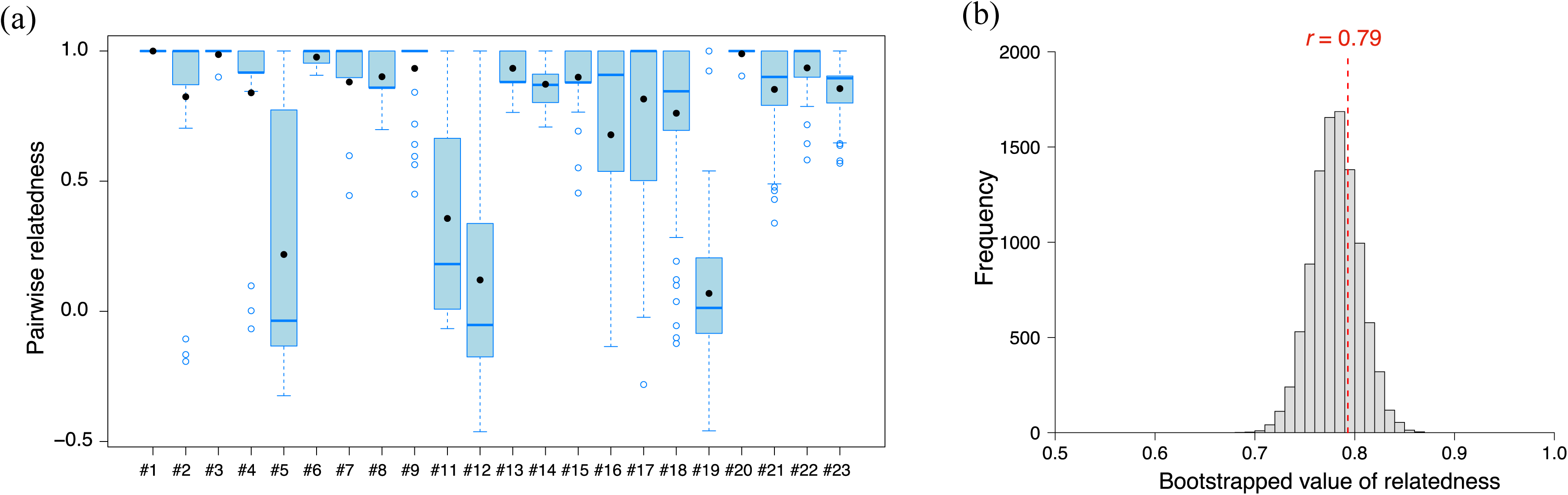
Genetic relatedness between head-butting aphids. (a) Within-colony pairwise genetic relatedness in *A. bambusae* (*N* = 22). Box plots show the range of values, the first and third quartiles and medians. Open circles show outliers and filled circles show means. (b) The distribution of pairwise relatedness within the colonies, derived from 10,000 bootstrapped values, with the red dotted line denoting the observed pairwise relatedness value (mean ± SD = 0.81 ± 0.10) for the 75 butting pairs.

Interestingly, 56 % (42/75) of the contests were performed by a clonal pair of aphids. Relatedness was not significantly associated with age difference, attacker age, or defender age or outcome of the fight (Table A2).

### Determinants of butting duration and outcome

The attacker won in 57% (63/111) of the contests, with no significant deviation from the expected value of 0.5 (χ^2^_1_ = 2.03, *P* = 0.15). In 60 out of 111 contests where age classes of the pair differed, older aphids had larger body size in all cases, and they were more likely to win (in 83% (50/60) of these contests, the older individual won: χ^2^_1_ = 26.7, *P* < 0.0001).

The GLMM analysis revealed that age class difference, whether attacker won the contest, and the age of defender were included in the final model (Table 2). Butting duration was significantly longer when age class differences were smaller (β ± SE = -0.26 ± 0.09, *Z* = -2.93, *P* = 0.003), when the attacker won (β ± SE = 0.48 ± 0.15, *Z* = 3.19, *P* = 0.001), and when the defender was older (β ± SE = 0.18 ± 0.08, *Z* = 2.28, *P* = 0.022, Table 2 and Fig. 4a). To investigate the influence of relatedness between the contestants on the duration of contest, we analysed 75 genotyped pairs engaged in contests. As in the comprehensive analysis that considered all individuals, we included age class difference, the contest outcome (whether the attacker won), and the age of defender in the final model (Table 2). Notably, relatedness was not a significant determinant in the duration of butting (β ± SE = -0.04 ± 0.25, *Z* = -0.16, *P* = 0.87). When categorizing the genotyped pairs into clonal and non-clonal pairs, the duration of butting was not significantly different between these groups (mean ± SD: 3.1 ± 2.6 min for clonal pairs; 2.5 ± 1.9 min for non-clonal pairs; Welch’s t-test, *t*_72.6_= 1.22, *P* = 0.22, Fig. 4b). The pairwise relatedness was not significantly different between contests won by the attacker (mean ± SD: 0.82 ± 0.35, *N* = 42) and those won by the defender (mean ± SD: 0.76 ± 0.34, *N* = 33; two-tailed bootstrap test, *P* = 0.50). These results suggest that the butting behaviour was not influenced by the genetic relatedness between the contestants.

**Figure 4.**
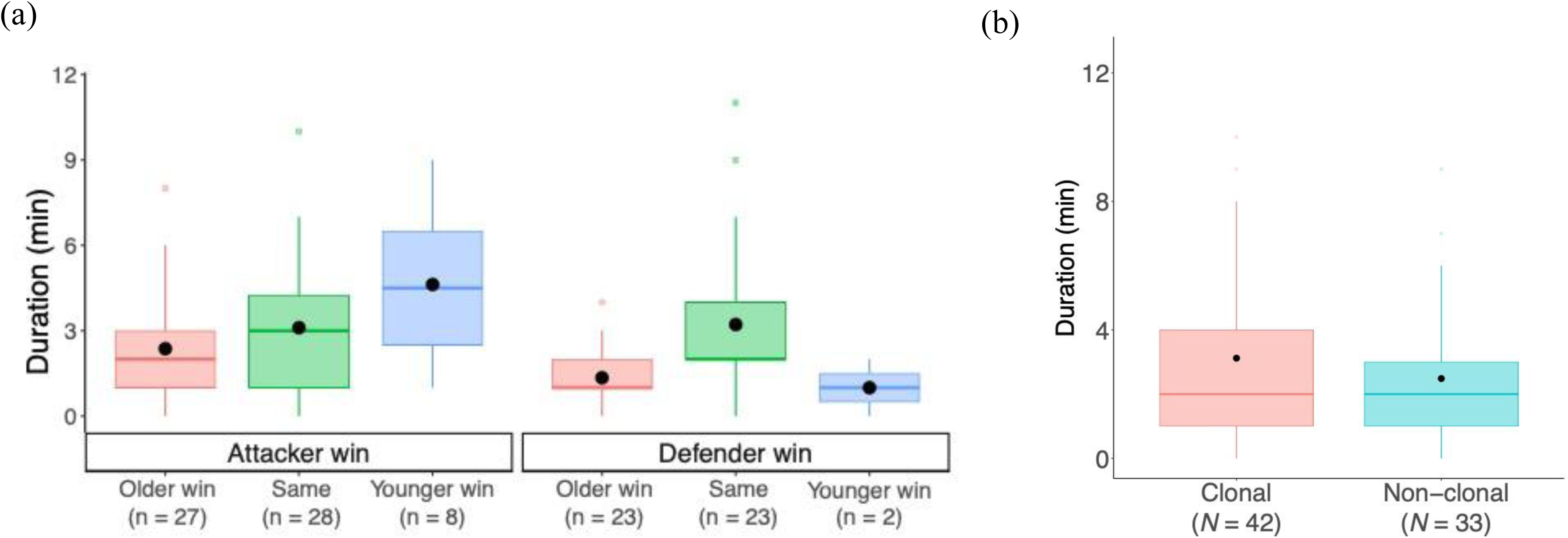
Determinants of the duration of contests. (a) Duration of contests won by attackers (left, *N* = 63) and defenders (right, *N* = 48). (b) Duration of contests between clonal (*N* = 41) and non-clonal (*N* = 34) pairs. Box plots show the range of values, the first and third quartiles and medians. Open circles show outliers and filled circles show means.

**Table 2.**
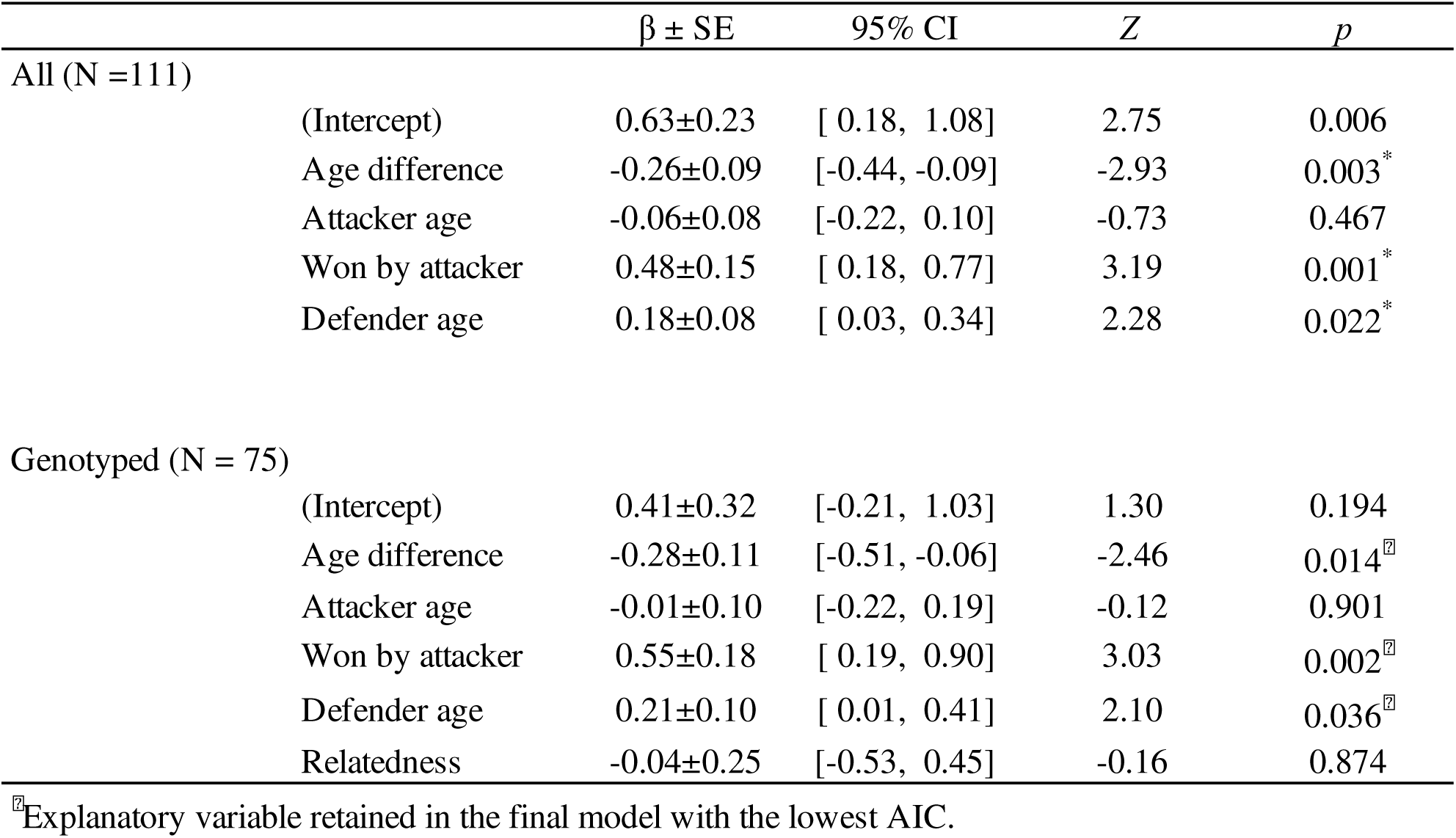
Relationships between butting duration and characteristics of competing aphids.

## Discussion

### What are the aphids fighting for?

These field observations provide the most complete account currently available of the duelling behaviour of a cerataphidine aphid and our novel measurements of genetic relatedness allow the adaptive value of this behaviour to be evaluated for the first time.

It is clear from previous work on *Astegopteryx* species, and this is probably true for most of the other cerataphidines, that the aphids on the secondary host are fighting for precise feeding sites (Aoki & Kurosu, 1985; Foster, 1996). Winning a feeding site by fighting can provide huge benefits in terms of the time saved in gaining access to a reliable source of food (Morris & Foster, 2008). The costs and benefits of fighting are best evaluated in terms of time: if the attacker wins, it gains the time which would otherwise have to be spent searching and inserting into a different feeding site, and the defender loses the time that it now needs to spend securing a new feeding site. Conversely, if the attacker loses, it then bears the time cost of finding a new feeding site; the winning defender bears few costs, since even during the fight she could continue to feed. Time not spent feeding really matters to aphids: because of the high nutritional requirements of rapidly developing embryos, they cannot live long without food.

### The ecology of fighting behaviour

Our observations are consistent with earlier studies on *Astegopteryx* aphids (Aoki & Kurosu, 1985; Foster, 1996), but are much more detailed and also more reliable since they were entirely carried out under field conditions. Laboratory observations on duelling aphids are compromised because the leaves dry out rapidly, which is one of the main reasons why aphids seek new feeding sites, thereby artificially increasing the frequency of fighting (Carlin et al., 1994; Foster, 1996). Unusually for a colony of feeding aphids, there is always a small but significant number of aphids (average 3%) moving about, attacking other aphids. The age distribution of both the attacking and defending aphids is highly non-random (Fig. 2). In particular, 1^st^ instars, although they are commonly seen to be duelling, in fact attack and defend significantly less frequently than their abundance would predict. These aphids must therefore be able to create – or perhaps find - their own feeding sites, since they are not born with access to the plant phloem (see also (Foster, 1996)). Adults are also significantly less likely to be involved in fights than their abundance would predict. This is presumably because other aphids usually choose not to attack them and they are therefore able to retain their initial feeding site. The majority of fights involve 2^nd^ and 3^rd^ instars, which are disproportionately likely to be seen acting as both defenders and attackers.

### The genetic context of fighting behaviour

Dyadic competitive interactions are usually studied in the context of the direct fitness outcomes for the contestants. We showed that older and larger aphids tended to win contests and that contest duration was influenced by the age difference between the contestants (Table 2). This finding is consistent with earlier studies on *Astegopteryx* aphids (Aoki & Kurosu, 1985; Foster, 1996), suggesting that the butting aphid pairs maximize their direct fitness benefits by assessing their opponent’s resource holding potential.

However, earlier work has shown that these aphids live in colonies consisting of a relatively small number of clones and average within-colony pairwise relatedness was 0.54 (Uematsu et al., 2023). It is therefore essential to consider not just the direct fitness of the contestants, but the inclusive fitness outcomes of the attacker and the defender in any contest. Our novel genetic analysis established that the attacking aphids, although presented with aphids of a huge range of potential relatedness values, choose to fight with aphids at random: they do not either favour or ignore aphids from the same clone (Fig. 3b). Nevertheless, the average pairwise relatedness of the contesting aphids was 0.79 and 56% of the contests *were* between aphids from the same clone.

What would be the relative inclusive fitness costs of winning in fights between clone- mates? Fighting itself does not seem to impose any detectable direct costs, such as wounds, on the contestants, and the duration of fighting in this study (Fig. 4) and on *A. minuta* (Foster, 1996) (mean: 2.7 min and 3.3 min, respectively) is very small compared to the time required to find and occupy a new feeding site. Therefore, the larger (that is older) attacker should win, since its reproductive value is higher than that of the younger defender: the time penalty of not gaining the feeding site (which could be very high) will impose a relatively higher fitness cost on the attacker, which might be close to reproducing, than on the younger defender. However, if the attacker is smaller, this same argument would imply that the defender should keep hold of its feeding site. This interpretation is consistent with earlier discussions that age or size can indicate future reproduction or reproductive value and can therefore influence altruistic behaviour between kin (Cant & Field, 2001; Rodrigues & Gardner, 2022).

### What are the aphids signalling to each other during a contest?

If this approach is correct, then a duel is not in fact a fight but rather an efficient, low-cost method that the aphids use to estimate each other’s relative reproductive value. Carlin et al. (Carlin et al., 1994)mainly focussed on the direct fitness value of fighting in their study, but they did suggest that the butting behaviour might also spread information through the colony about the state of desiccation of the host plant. It is empirically very difficult to distinguish between cooperation and aggression in dyadic contexts. The basic difference is that aggressive fights are usually structured around relative resource holding potential, whereas cooperative encounters are about relative reproductive value. In both contexts, but for different reasons, the contests usually include an element of assessment; the older and larger contestant generally wins; and fights last longer when the opponents are more evenly matched in age. In both cases, very young animals rarely attack adults, since they can usually detect, without the need for contact, that these larger individuals are not worth attacking. It seems reasonable to conclude, therefore, that the aphid encounters are cooperative, since the interactions in *Astegopteryx* never involve injury or excessive time-costs. Aphids in this clade are nevertheless perfectly capable of evolving damaging weaponry and longer fights (see for example (Aoki & Kurosu, 2010; Howard et al., 1998)).

How might aphids assess each other’s reproductive value? It seems likely that they do this simply by “measuring” each other’s size, which is in general a reliable indicator of age (Foster, 1996). Clasping will show the attacker how wide its opponent is and doing a handstand would be a good way for a defender to show long and heavy it is. Head-butting another aphid might also give some idea of its mass.

### Why does relatedness not affect the occurrence, duration or outcome of contests?

Relatedness does not affect which aphids the attackers chose to attack (Fig 3b), or the duration or outcome of a contest (Table 2, Fig 4b). This finding corroborates a previous study on another cerataphidine, *Ceratovacuna japonica* (Carlin et al., 1994), and supports the absence of differences in inter- and intra-species head-butting duration in a mixed colony of two *Astegopteryx* species (Aoki & Kurosu, 1985). In fact, there is very little evidence that aphids can discriminate between clone and non-clone mates, in any context, including defensive behaviour (Miller III., 1998; Shibao, 1999); and mating (Foster & Benton, 1992) (but see also Li & Akimoto, 2021).

Previous studies of duelling in *Astegopteryx* have assumed that the aphids would benefit from evicting non-clone mates from their feeding sites. But if our view that the contests are in fact cooperative is correct, then we would expect natural selection to favour contests with clone-mates. It is difficult to envisage how an aphid might be selected to behave differentially when encountering a clone vs a non-clone mate, since this differential behaviour has never been detected. But any scenario would assume that the fights would be much costlier, perhaps both in time and in injuries: the defender would gain no indirect fitness advantage by withdrawing and the attacker would not incur any costs by forcing a non-clone mate from its feeding site. Therefore, if clones were identifiable by genetic tags, when an aphid of an alien clone joins an existing pure clone, it will almost always be involved in contests, either as an attacker or a defender, with aphids of a different clone. These contests could be more costly than the much commoner cooperative interactions between aphids of the pure clone: following a version of Crozier’s Paradox, this invading clone and its tag, and as result genetic tag diversity, would be eliminated (Crozier, 1986; Field et al., 2018). One eriosomatine aphid (*Pemphigus obesinymphae*) is able to detect when they are in a colony of non-clone mates, but only if they have migrated to a distinct aphid colony (Abbot et al., 2001). It would be worth testing whether a *Astegopteryx* aphids that have migrated to totally new colonies behave as though they are surrounded by non-clone mates.

### The evolution of duelling and of the soldier caste in aphids

It has been suggested that the duelling behaviour of cerataphidine aphids might have influenced the evolution of the anti-predator fighting behaviour of the horned sterile 1^st^ instar soldiers on the secondary host (Aoki, 1987; Aoki & Kurosu, 1985; Foster, 1996; Stern & Foster, 1996). Our observations show that, although the 1^st^ instars are significantly less likely to attack than predicted from their abundance in a colony, they nevertheless are frequently observed to attack and defeat 1^st^-instar defenders. Our data also provides a more general context for why altruistic behaviour might have become focussed on the 1^st^ instars: they have the smallest reproductive value and therefore, if duelling is cooperative, should be more likely to yield their feeding sites to older aphids, promoting larger differences in future reproductive success, consistent with theoretical prediction (Johnstone, 2008).

Duelling behaviour has been recorded in all the examined cerataphidine species (Aoki & Kurosu, 2010). However, its intensity seems to vary in different species, which could give us clues about how the behaviour might have evolved. There appears to be a spectrum of intensity from *Cerataphis brasiliensis*, where duelling is often prolonged (> 14 min), weaponised with long sharp horns, and potentially injurious (Aoki, 1987; Howard et al., 1998), through the apparently cooperative behaviour described in *Astegopteryx*, to the mild fighting often observed in *Pseudoregma* and *Ceratovacuna* (Aoki & Kurosu, 2010). It would be interesting to investigate the intensity of fighting behaviour in relation to genetic relatedness in colonies of other cerataphidine aphids in the light of a rigorous phylogenetic analysis.

## Data Availability

The data and R code for this study are available as Supplementary material.

## Acknowledgements

This work was supported by supported by the Sumitomo Foundation (130972) and JSPS Overseas Research Fellowships (24-596) to KU. We thank Chun-I Chiu, Bao-Cheng Lai, Yi- Chuan Lee, and Chang-Ti Tang for help with fieldwork.

---

**Supplementary figure 1.**
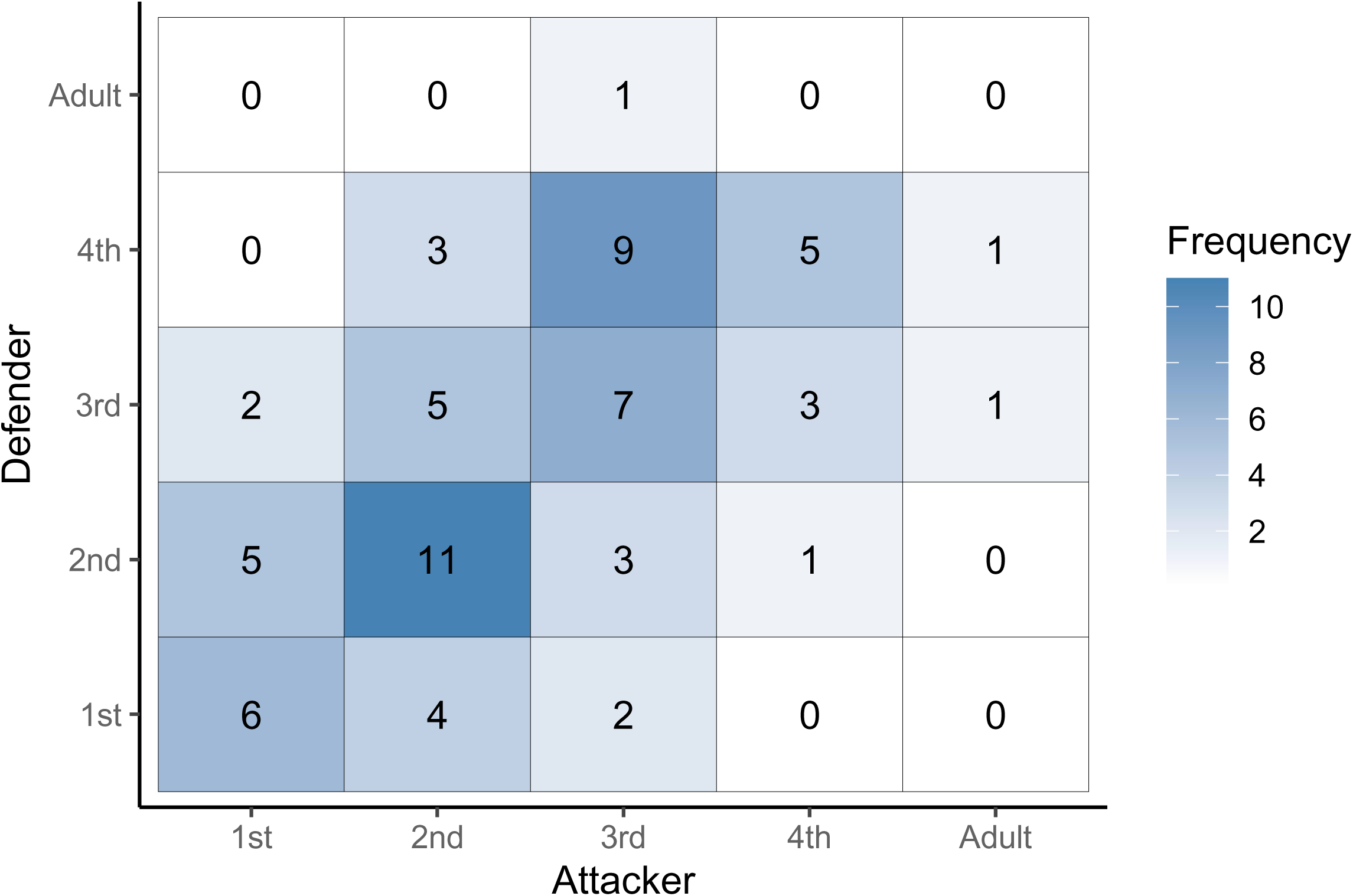
Pairwise frequency matrix in the age of attackers and defenders. Color density indicate the frequency in the combination of the five age categories (1^st^ to 4^th^ instar and adult).

Supplementary Video S1. Head-butting behaviour of *Astegopteryx bambusae*.

**Table A1.** The success rates of attacks on different ages.

| Age of defender | Number of attackers |  | % success of attacks by |  |  |
| --- | --- | --- | --- | --- | --- |
|  | Younger | Older | Younger | Older | Overall* |
| 1st | 0 | 15 | na | 100% (15/15) | 100% (15/15) |
| 2nd | 7 | 7 | 29% (2/7) | 100% (7/7) | 64% (9/14) |
| 3rd | 7 | 6 | 29% (2/7) | 83% (5/6) | 54% (7/13) |
| 4th | 13 | 1 | 23% (3/13) | 0% (0/1) | 21% (3/14) |
| Adult | 4 | 0 | 25% (1/4) | na | 25% (1/4) |

\* $P < 0.0001$ (Cochran-Armitage trend test, $\chi_1 = 19.2$ )

**Table A2.** Relationships between relatedness and characteristics.

| | $\beta \pm \text{SE}$ | 95% CI | $t_{68}$ | $P$ |
| --- | --- | --- | --- | --- |
| (Intercept) | 0.62±0.12 | [ 0.37, 0.86] | 5.03 | < 0.001 |
| Age difference | 0.03±0.05 | [-0.06, 0.12] | 0.67 | 0.505 |
| Attacker age | 0.07±0.04 | [-0.01, 0.15] | 1.72 | 0.09 |
| Defender age | 0±0.04 | [ -0.08, 0.08] | -0.09 | 0.932 |
| Won by attacker | -0.01±0.08 | [ -0.17, 0.15] | -0.08 | 0.938 |

## References

Abbot, P., Withgott, J. H., & Moran, N. A. (2001). Genetic conflict and conditional altruism in social aphid colonies. Proceedings of the National Academy of Sciences of the United States of America, 98(21), 12068–12071.

Amos, W., Nichols, H. J., Churchyard, T., & Brooke, M. de L. (2016). Rat eradication comes within a whisker! A case study of a failed project from the South Pacific. R. Soc. Open Science, 3(160110), 1–10. 10.1098/rsos.160110

Aoki, S. (1987). Evolution of sterlile soldiers in aphids. In Y. Ito et al (Ed.), Animal Societies: Theories and Facts (pp. 53–65). Japan Science Society Press.

Aoki, S., & Kurosu, U. (1985). An aphid species doing a headstand: Butting behavior of *Astegopteryx bambucifoliae* (Homoptera: Aphidoidea). Journal of Ethology, 3(2), 83–87. 10.1007/BF02350297

Aoki, S., & Kurosu, U. (2010). A Review of the Biology of Cerataphidini (Hemiptera, Aphididae, Hormaphidinae), Focusing Mainly on Their Life Cycles, Gall Formation, and Soldiers. Psyche: A Journal of Entomology, 2010, 1–34. 10.1155/2010/380351

Bates, D., Mächler, M., Bolker, B., & Walker, S. (2015). Fitting Linear Mixed-Effects Models Using lme4. Journal of Statistical Software, 67(1). 10.18637/jss.v067.i01

Bell, G. (1982). The masterpiece of nature: the evolution and genetics of sexuality. CUP Archive.

Cant, M. A., & Field, J. (2001). Helping effort and future fitness in cooperative animal societies. Proceedings of the Royal Society of London. Series B: Biological Sciences, 268(1479), 1959–1964. 10.1098/rspb.2001.1754

Carlin, N., Gladstein, D., Berry, A., & Pierce, N. (1994). Absence of kin discrimination behavior in a soldier-producing aphid, Ceratovacuna japonica (Hemiptera: Pemphigidae; Cerataphidini). J. New York Entomol. Soc., 102(3), 287–298. http://www.jstor.org/stable/10.2307/25010090

Croft, D. P., Weiss, M. N., Nielsen, M. L. K., Grimes, C., Cant, M. A., Ellis, S., Franks, D. W., & Johnstone, R. A. (2021). Kinship dynamics: Patterns and consequences of changes in local relatedness. In Proceedings of the Royal Society B: Biological Sciences (Vol. 288, Issue 1957). Royal Society Publishing. 10.1098/rspb.2021.1129

Crozier, R. H. (1986). Genetic clonal recognition abilities in marine invertebrates must be maintained by selection for something else. Evolution, 40(5), 1100–1101. 10.1111/J.1558-5646.1986.TB00578.X

Field, J., Accleton, C., & Foster, W. A. (2018). Crozier’s Effect and the Acceptance of Intraspecific Brood Parasites. Current Biology, 28(20), 3267–3272.e3. 10.1016/J.CUB.2018.08.014

Foster, W. A. (1996). Duelling aphids: Intraspecific fighting in *Astegopteryx minuta* (Homoptera: Hormaphididae). Animal Behaviour, 51, 645–655. 10.1006/anbe.1996.0069

Foster, W. A., & Benton, T. G. (1992). Sex ratio, local mate competition and mating behaviour in the aphid *Pemphigus spyrothecae*. Behavioral Ecology and Sociobiology, 30(5), 297–307. 10.1007/BF00170595

Green, P. A., Briffa, M., & Cant, M. A. (2021). Assessment during Intergroup Contests. In Trends in Ecology and Evolution (Vol. 36, Issue 2, pp. 139–150). Elsevier Ltd. 10.1016/j.tree.2020.09.007

Hardy, I. C. W., & Briffa, M. (2013). Animal contests. Cambridge University Press.

Hardy, I. C. W., & Mesterton-Gibbons, M. (2023). The evolutionarily stable strategy, animal contests, parasitoids, pest control and sociality. In Philosophical Transactions of the Royal Society B: Biological Sciences (Vol. 378, Issue 1876). Royal Society Publishing. 10.1098/rstb.2021.0498

Hasegawa, M., & Kutsukake, N. (2019). Kin selection and reproductive value in social mammals. Journal of Ethology, 37(2), 139–150. 10.1007/s10164-019-00586-6

Howard, F. W., Halbert, S., & Giblin-davis, R. (1998). Intraspecific Dueling in Palm Aphids, *Cerataphis brasiliensis* (Homoptera: Hormaphididae). The Florida Entomologist, 81(4), 552. 10.2307/3495956

Hughes, R. N. (1989). Functional biology of clonal animals. Springer Science & Business Media.

Johnstone, R. A. (2008). Kin selection, local competition, and reproductive skew. Evolution, 62(10), 2592–2599. 10.1111/j.1558-5646.2008.00480.x

Kokko, H. (2013). Dyadic contests: modelling fights between two individuals. In Animal Contests (pp. 5–32). Cambridge University Press. 10.1017/CBO9781139051248.004

Konovalov, D. A., Manning, C., & Henshaw, M. T. (2004). KINGROUP: A program for pedigree relationship reconstruction and kin group assignments using genetic markers. Molecular Ecology Notes, 4(4), 779–782. 10.1111/j.1471-8286.2004.00796.x

Li, Q., Yao, J., Zeng, L., Lin, X., & Huang, X. (2019). Molecular and morphological evidence for the identity of two nominal species of *Astegopteryx* (Hemiptera, Aphididae, Hormaphidinae). ZooKeys, 833, 59–74. 10.3897/zookeys.833.30592

Li, Y., & Akimoto, S. (2021). Self and non-self recognition affects clonal reproduction and competition in the pea aphid. Proceedings of the Royal Society B: Biological Sciences, 288(1953). 10.1098/rspb.2021.0787

Miller III., D. G. (1998). Consequences of communal gall occupation and a test for kin discrimination in the aphid *Tamalia coweni* (Cockerell) (Homoptera: Aphididae). Behavioral Ecology and Sociobiology, 43(2), 95–103. 10.1007/s002650050471

Morris, G., & Foster, W. A. (2008). Duelling aphids: electrical penetration graphs reveal the value of fighting for a feeding site. Journal of Experimental Biology, 211(9), 1490– 1494. 10.1242/jeb.012120

Parker, G. A. (1974). Assessment strategy and the evolution of fighting behaviour. Journal of Theoretical Biology, 47(1), 223–243. 10.1016/0022-5193(74)90111-8

Queller, D. C., & Goodnight, K. F. (1989). Estimating relatedness using genetic markers. Evolution, 43(2), 258. 10.2307/2409206

R Core Team, A., & Team, R. C. (2022). R: A language and environment for statistical computing. *R Foundation for Statistical Computing*, *Vienna*, *Austria*.

Rodrigues, A. M. M., & Gardner, A. (2022). Reproductive value and the evolution of altruism. Trends in Ecology & Evolution, 37(4), 346–358. 10.1016/j.tree.2021.11.007

Shibao, H. (1999). Lack of kin discrimination in the eusocial aphid *Pseudoregma bambucicola* (Homoptera[: Aphididae). Journal of Ethology, 17(1), 17–24.

Stern, D. L., & Foster, W. A. (1996). The evolution of soldiers in aphids. Biological Reviews of the Cambridge Philosophical Society, 71(1), 27–79.

Taborsky, M., Cant, M. A., & Komdeur, J. (2021). The Evolution of Social Behaviour. Cambridge University Press.

Uematsu, K., Shimada, M., & Shibao, H. (2013). Juveniles and the elderly defend, the middle-aged escape: division of labour in a social aphid. Biology Letters, 9(2), 20121053–20121053. 10.1098/rsbl.2012.1053

Uematsu, K., Yang, M.-M., Amos, W., & Foster, W. A. (2023). Eusocial evolution without a nest: kin structure of social aphids forming open colonies on bamboo. Behavioral Ecology and Sociobiology, 77(3), 38. 10.1007/s00265-023-03315-9

Zuur, A. F., Ieno, E. N., & Elphick, C. S. (2010). A protocol for data exploration to avoid common statistical problems. Methods in Ecology and Evolution, 1(1), 3–14. 10.1111/j.2041-210X.2009.00001.x

