## Supplementary data and R code for "The kin-selected context of duelling in horned aphids: cooperation or conflict?": age_bootstrap_test.R

```
library(tidyverse)

data <- read.csv('age_bootstrap_test.csv')

s <- numeric()
sa <- numeric()
sd <- numeric()
for(l in 1:10000){
j <- numeric()
for(k in 13:23){
eval(parse(text =paste0("a <- subset(data,colony=='",k,"')")))
b <- c(rep(1,rep(a[,2])),rep(2,rep(a[,3])),rep(3,rep(a[,4])),rep(4,rep(a[,5])),rep(5,rep(a[,6])))
for (i in 1:a[,7]){
d <- sample(b,2,replace=TRUE)
j <- rbind(j,d)
}
}
r <- numeric()
r[1]<-sum(j[,1]==1)+sum(j[,2]==1)
r[2]<-sum(j[,1]==2)+sum(j[,2]==2)
r[3]<-sum(j[,1]==3)+sum(j[,2]==3)
r[4]<-sum(j[,1]==4)+sum(j[,2]==4)
r[5]<-sum(j[,1]==5)+sum(j[,2]==5)
s <- rbind(s,r)

ra <- numeric()
ra[1]<-sum(j[,1]==1)
ra[2]<-sum(j[,1]==2)
ra[3]<-sum(j[,1]==3)
ra[4]<-sum(j[,1]==4)
ra[5]<-sum(j[,1]==5)
sa <- rbind(sa,ra)

rd <- numeric()
rd[1]<-sum(j[,2]==1)
rd[2]<-sum(j[,2]==2)
rd[3]<-sum(j[,2]==3)
rd[4]<-sum(j[,2]==4)
rd[5]<-sum(j[,2]==5)
sd <- rbind(sd,rd)

}
summary(s)
summary(sa)
summary(sd)
#top 5%
x1 <- apply(s,2,function(x){
sort(x, decreasing=TRUE)[500]
})
xa1 <- apply(sa,2,function(x){
sort(x, decreasing=TRUE)[500]
})
xd1 <- apply(sd,2,function(x){
sort(x, decreasing=TRUE)[500]
})
#bottom 5%
x2 <- apply(s,2,function(x){
sort(x, decreasing=TRUE)[9501]
})
xa2 <- apply(sa,2,function(x){
sort(x, decreasing=TRUE)[9501]
})
xd2 <- apply(sd,2,function(x){
sort(x, decreasing=TRUE)[9501]
})

#median
xm <- apply(s,2,function(x){
sort(x, decreasing=TRUE)[5000]
})
xam <- apply(sa,2,function(x){
sort(x, decreasing=TRUE)[5000]
})
xdm <- apply(sd,2,function(x){
sort(x, decreasing=TRUE)[5000]
})


#observed data
obs_attacker <- c(13,23,22,9,2)
obs_defender <- c(12,20,18,18,1)
obs_pair <- obs_attacker+obs_defender
x12 <-as.data.frame(cbind(x1,x2,xm,obs_pair))
xad <-as.data.frame(cbind(xa1,xa2,xam,obs_attacker,obs_defender))


colnames(x12) <- c("top","bottom","median","obs")
x12$age <- c("1st","2nd","3rd","4th","adult")

colnames(xad) <- c("top","bottom","median","obs_a","obs_d")
xad$age <- c("1st","2nd","3rd","4th","adult")

ggplot(data=x12) + aes(x=age)+
  geom_errorbar(aes(ymin=bottom,ymax=top),linetype=1,color="black") +
  geom_point(aes(y=obs),size=3,color="red")+
  geom_point(aes(y=median),shape=4,size=2,color="black")+
  theme_classic()+ylab("Frequency")+xlab("Age")
ggsave(file="age_bootstrap_combined.pdf")
  

ggplot(data=xad) + aes(x=age)+
 geom_errorbar(aes(ymin=bottom,ymax=top),linetype=1,color="black") +
  geom_point(aes(y=obs_a),size=3,color="red")+
  geom_point(aes(y=obs_d),size=3,color="blue")+
  geom_point(aes(y=median),shape=4,size=2,color="black")+
  theme_classic()+ylab("Frequency")+xlab("Age")
ggsave(file="age_bootstrap_attack-defend.pdf")
  

#real data
hbpairs_field <- read.csv("hbpairs_field.csv")
rev <- subset(hbpairs_field,hbpairs_field$group >= 13 & hbpairs_field$group <= 23)
table(rev$age_attacker, rev$age_defender)
j2 <- data.frame(rev$age_attacker,rev$age_defender)
q <- numeric()
q[1]<-sum(j2[,1]==1)+sum(j2[,2]==1)
q[2]<-sum(j2[,1]==2)+sum(j2[,2]==2)
q[3]<-sum(j2[,1]==3)+sum(j2[,2]==3)
q[4]<-sum(j2[,1]==4)+sum(j2[,2]==4)
q[5]<-sum(j2[,1]==5)+sum(j2[,2]==5)
q

qa <- numeric()
qa[1]<-sum(j2[,1]==1)
qa[2]<-sum(j2[,1]==2)
qa[3]<-sum(j2[,1]==3)
qa[4]<-sum(j2[,1]==4)
qa[5]<-sum(j2[,1]==5)
qa

qd <- numeric()
qd[1]<-sum(j2[,2]==1)
qd[2]<-sum(j2[,2]==2)
qd[3]<-sum(j2[,2]==3)
qd[4]<-sum(j2[,2]==4)
qd[5]<-sum(j2[,2]==5)
qd


tab <- as.data.frame(table(rev$age_attacker, rev$age_defender))
colnames(tab) <- c("Attacker_age","Defender_age","Frequency")
instar <- c("1st","2nd","3rd","4th","Adult")
ggplot(tab,aes(Attacker_age,Defender_age),fill = Frequency) + geom_tile(aes(fill=Frequency),colour="black") + geom_text(aes(label=Frequency),colour = "black")+labs(x= "Attacker",y = "Defender") +theme_classic()+ scale_fill_gradient(low="white",high="steelblue",breaks=c(2, 4, 6, 8, 10), limits=c(0, 11))+scale_x_discrete(labels = instar)+scale_y_discrete(labels = instar)
ggsave(file="age_table.pdf")
```
