## Supplementary data and R code for "The kin-selected context of duelling in horned aphids: cooperation or conflict?": headbutt_analysis.R

```
options(max.print=999999) 
library('tidyverse')
library('dplyr')
library('lme4')
library('lmerTest')
library('car')
library('parameters')

data <- read.csv("data.csv")
data$relatedness <- as.numeric(data$relatedness)
data$num <- as.numeric(data$num)
summary(data)
hbpairs <- subset(data, data$group==data$group_pair & data$headbutt=="Y" & data$headbutt_pair=="Y" & data$num==data$num_pair & data$outcome=="W")
summary(hbpairs)
hbpairs_clone <- subset(hbpairs, hbpairs$relatedness==1)
hbpairs_clone


#frequency-colony size
colony <- read.csv("Colony sizes.csv")
summary(colony$number.of.aphids)
sd(colony$number.of.aphids)/sqrt(length(colony$number.of.aphids))
testResult <- cor.test(colony$number.of.aphids, colony$frequency)
testResult ##t = 0.76897, df = 21, p-value = 0.4505
summary(colony$attacker_per_min)
sd(colony$attacker_per_min)
sd(colony$attacker_per_min)/sqrt(length(colony$attacker_per_min))
colony$attacker_per_hour <- (colony$frequency*2)/colony$number.of.aphids
summary(colony$attacker_per_hour)
sd(colony$attacker_per_hour)
sd(colony$attacker_per_hour)/sqrt(length(colony$attacker_per_hour))

hbpairs_field <- read.csv("hbpairs_field.csv")

#defender instars and success rate
def_1st_old <- subset(hbpairs_field,hbpairs_field$age_defender =="1" & hbpairs_field$age_diff_noabs > 0 )
def_2nd_yon <- subset(hbpairs_field,hbpairs_field$age_defender =="2" & hbpairs_field$age_diff_noabs < 0 )
def_2nd_old <- subset(hbpairs_field,hbpairs_field$age_defender =="2" & hbpairs_field$age_diff_noabs > 0 )
def_3rd_yon <- subset(hbpairs_field,hbpairs_field$age_defender =="3" & hbpairs_field$age_diff_noabs < 0 )
def_3rd_old <- subset(hbpairs_field,hbpairs_field$age_defender =="3" & hbpairs_field$age_diff_noabs > 0 )
def_4th_yon <- subset(hbpairs_field,hbpairs_field$age_defender =="4" & hbpairs_field$age_diff_noabs < 0 )
def_4th_old <- subset(hbpairs_field,hbpairs_field$age_defender =="4" & hbpairs_field$age_diff_noabs > 0 )
def_ad_yon <- subset(hbpairs_field,hbpairs_field$age_defender =="5" & hbpairs_field$age_diff_noabs < 0 )
#Cochran-Armitage test
success <- c(15,9,7,3,1)
total <- c(15,14,13,14,4)
prop.trend.test(success, total)
#X-squared = 19.215, df = 1, p-value = 1.168e-05

#clones or non-clonal
clones <- subset(hbpairs_field, relatedness==1)
nonclones <- subset(hbpairs_field, relatedness<1)
summary(clones)
summary(nonclones)
clones$clonality <- as.factor("Clonal")
nonclones$clonality <- as.factor("Non-clonal")

genotyped <- rbind(clones,nonclones)

#Relatedness explanatory
rel1 <- lmer(relatedness~age_diff+age_attacker+age_defender+won.by.attacker+(1|group), data = genotyped) 
summary(rel1)
model_parameters_rel1 <- model_parameters(rel1)
model_parameters_rel1

#winnerの検定 attacker > defender? 
hbpairs_field$won.by.attacker <- as.factor(hbpairs_field$won.by.attacker)
summary(hbpairs_field$won.by.attacker) #attacker:63, defender:48
won.by.attacker <- c (63,48)
prob <- c(0.5,0.5)
testResult <- chisq.test(x = won.by.attacker, p = prob)
testResult
# chi-squared = 2.027, P = 0.15

#won.by.attacker larger > smaller test
attacker_lw <- subset(hbpairs_field,hbpairs_field$won.by.attacker =="1" & hbpairs_field$age_diff_noabs > 0 )
attacker_same <- subset(hbpairs_field,hbpairs_field$won.by.attacker =="1" & hbpairs_field$age_diff_noabs == 0 )
attacker_sw <- subset(hbpairs_field,hbpairs_field$won.by.attacker =="1" & hbpairs_field$age_diff_noabs < 0 )
defender_lw <- subset(hbpairs_field,hbpairs_field$won.by.attacker =="0" & hbpairs_field$age_diff_noabs < 0 )
defender_same <- subset(hbpairs_field,hbpairs_field$won.by.attacker =="0" & hbpairs_field$age_diff_noabs == 0 )
defender_sw <- subset(hbpairs_field,hbpairs_field$won.by.attacker =="0" & hbpairs_field$age_diff_noabs > 0 )
equal <- subset(hbpairs_field,hbpairs_field$age_diff_noabs == 0 )

large_win <- nrow(attacker_lw)+nrow(defender_lw)
small_win <- nrow(attacker_sw)+nrow(defender_sw)
winner <- c (large_win,small_win)
prob <- c(0.5,0.5)
testResult <- chisq.test(x = winner, p = prob)
testResult
# chi-squared =26.667, P < 0.0001


#duration 
summary(attacker_lw$duration)
summary(attacker_same$duration)
summary(attacker_sw$duration)
summary(defender_lw$duration)
summary(defender_same$duration)
summary(defender_sw$duration)

#boxplot of duration
attacker_lw$diff <- "Older win"
attacker_lw$winner <- "Attacker win"
attacker_same$diff <- "Same"
attacker_same$winner <- "Attacker win"
attacker_sw$diff <- "Younger win"
attacker_sw$winner <- "Attacker win"

defender_lw$diff <- "Older win"
defender_same$diff <- "Same"
defender_sw$diff <- "Younger win"
defender_lw$winner <- "Defender win"
defender_same$winner <- "Defender win"
defender_sw$winner <- "Defender win"
data <- rbind(attacker_lw,attacker_same,attacker_sw,defender_lw,defender_same,defender_sw)

Boxplot <- ggplot(data, aes(x = diff, y = duration, colour = diff, fill = diff)) +geom_boxplot(alpha = 0.40) + facet_wrap(~winner, ncol = 3, switch = "x",scales="fixed") + coord_cartesian(ylim = c(0,11.5)) +theme_classic() +theme(axis.text.x  = element_text(size=14), axis.text.y = element_text(size=14), strip.text.x = element_text(size =16)) + theme(axis.title.x = element_text(size=14), axis.title.y = element_text(size=16), plot.title=element_text(size=0)) + xlab("") + ylab("Duration (min)") + theme(legend.position = "none")+ stat_summary(fun.y=mean, geom="point", shape=20, size=5, color="black", fill="black")
Boxplot
ggsave(file="duration_outcomes.pdf",width = 8, height = 4)


#welch's t test , clonal vs non-clonal #figure 4b
t.test(clones$duration,nonclones$duration,var.equal=FALSE)
# calculate standard error
print(sd(clones$duration))
print(sd(nonclones$duration))


#attacker win or defender win
attackerwin <- subset(hbpairs_field, hbpairs_field$won.by.attacker =="1")
defenderwin <- subset(hbpairs_field, hbpairs_field$won.by.attacker =="0")
summary(attackerwin)
summary(defenderwin)


#boxplot + dotplot-relatedness
#duration- clonality
genotyped$clonality <- as.factor(genotyped$clonality)
p <- ggplot(genotyped, aes_string(x ="clonality",  y= "duration" ,colour = "clonality", fill = "clonality"))
p <- p + ylim(c(0,12.5))+ geom_boxplot(alpha=0.4, outlier.alpha = 0.2, outlier.fill="white") +theme_classic() +theme(axis.text.x  = element_text(size=30), axis.text.y = element_text(size=30), strip.text.x = element_text(size =30)) + theme(axis.title.x = element_text(size=30), axis.title.y = element_text(size=30), plot.title=element_text(size=0)) + xlab("") + ylab("Duration (min)") + theme(legend.position = "none")+ stat_summary(fun.y=mean, geom="point", shape=20, size=5, color="black", fill="black")
plot(p)
ggsave("clonality_duration.pdf")


# glmm
#all pairs
res <- glmer(duration~age_diff+age_attacker+won.by.attacker+age_defender+(1|group), data = hbpairs_field,family="poisson")
summary(res)#AIC:455.2
model_parameters <- model_parameters(res)
model_parameters


#genotyped pairsonly
hbpairs_field3 <- subset(hbpairs_field, relatedness != "NA")
res_r <- glmer(duration~age_diff+age_attacker+won.by.attacker+age_defender+relatedness+(1|group), data = hbpairs_field3,family="poisson")#AIC:316.4
summary(res_r)
model_parameters_r <- model_parameters(res_r)
model_parameters_r

# vif
vif(res)
vif(res_r)

#model selection

res0 <- glmer(duration~age_diff+won.by.attacker+age_attacker+(1|group), data = hbpairs_field,family="poisson")
summary(res0)#AIC:458.5
vif(res0)

res1 <- glmer(duration~age_diff+won.by.attacker+age_defender+(1|group), data = hbpairs_field,family="poisson")
summary(res1)#AIC:453.8 #lowest
vif(res1)

res2 <- glmer(duration~age_diff+won.by.attacker+(1|group), data = hbpairs_field,family="poisson")
summary(res2)#AIC:457.0
vif(res2)

res3 <- glmer(duration~won.by.attacker+age_defender+(1|group), data = hbpairs_field,family="poisson")
summary(res3)#AIC:460.2
vif(res3)

res4 <- glmer(duration~age_diff+age_defender+(1|group), data = hbpairs_field,family="poisson")
summary(res4) #AIC:461.5
vif(res4)

## res1 has lowest AIC

##
##only genotyped pairs - table 2 lower part


res_genotyped0 <- glmer(duration~age_diff+age_attacker+won.by.attacker+age_defender+relatedness+(1|group), data = hbpairs_field3,family="poisson")
summary(res_genotyped0)#AIC:316.5

res_genotyped1 <- glmer(duration~age_diff+age_attacker+won.by.attacker+relatedness+(1|group), data = hbpairs_field3,family="poisson") 
summary(res_genotyped1)#AIC:319.0
vif(res_genotyped1)

res_genotyped2 <- glmer(duration~age_diff+age_defender+won.by.attacker+relatedness+(1|group), data = hbpairs_field3,family="poisson") 
summary(res_genotyped2)#AIC:314.5
vif(res_genotyped2)

res_genotyped3 <- glmer(duration~age_diff+age_defender+won.by.attacker+(1|group), data = hbpairs_field3,family="poisson") 
summary(res_genotyped3)#AIC:312.5 #lowest
vif(res_genotyped3)

res_genotyped4 <- glmer(duration~age_diff+won.by.attacker+(1|group), data = hbpairs_field3,family="poisson") 
summary(res_genotyped4)#AIC:317.4
vif(res_genotyped4)

res_genotyped5 <- glmer(duration~age_attacker+won.by.attacker+(1|group), data = hbpairs_field3,family="poisson") 
summary(res_genotyped5)#AIC:318.4
vif(res_genotyped5)

res_genotyped6 <- glmer(duration~age_diff+age_attacker+(1|group), data = hbpairs_field3,family="poisson") 
summary(res_genotyped6)#AIC:320.6

## res_genotyped3 has lowest AIC
```
