## Supplementary data and R code for "The kin-selected context of duelling in horned aphids: cooperation or conflict?": relatedness_tests_for_deposit.R

```
options(max.print=999999) 

relatedness<- read.csv("Group_results_rev.csv",header=TRUE)
relatedness$r <- as.numeric(relatedness$r)


group1 <- subset (relatedness, Group=="1" & PairGroup=="1")
group2 <- subset (relatedness, Group=="2" & PairGroup=="2")
group3 <- subset (relatedness, Group=="3" & PairGroup=="3")
group4 <- subset (relatedness, Group=="4" & PairGroup=="4")
group5 <- subset (relatedness, Group=="5" & PairGroup=="5")
group6 <- subset (relatedness, Group=="6" & PairGroup=="6")
group7 <- subset (relatedness, Group=="7" & PairGroup=="7")
group8 <- subset (relatedness, Group=="8" & PairGroup=="8")
group9 <- subset (relatedness, Group=="9" & PairGroup=="9")
group10 <- subset (relatedness, Group=="10" & PairGroup=="10")
group11 <- subset (relatedness, Group=="11" & PairGroup=="11")
group12 <- subset (relatedness, Group=="12" & PairGroup=="12")
group13 <- subset (relatedness, Group=="13" & PairGroup=="13")
group14 <- subset (relatedness, Group=="14" & PairGroup=="14")
group15 <- subset (relatedness, Group=="15" & PairGroup=="15")
group16 <- subset (relatedness, Group=="16" & PairGroup=="16")
group17 <- subset (relatedness, Group=="17" & PairGroup=="17")
group18 <- subset (relatedness, Group=="18" & PairGroup=="18")
group19 <- subset (relatedness, Group=="19" & PairGroup=="19")
group20 <- subset (relatedness, Group=="20" & PairGroup=="20")
group21 <- subset (relatedness, Group=="21" & PairGroup=="21")
group22 <- subset (relatedness, Group=="22" & PairGroup=="22")
group23 <- subset (relatedness, Group=="23" & PairGroup=="23")

#function for removing pairwise duplicates
remov_pwisedup <- function(x) {
indx <- !duplicated(t(apply(x[,c(1,3)], 1, sort))) # finds non - duplicates in sorted rows
return(x[indx, ])
}

#removing duplicates
for (i in c(1:9,11:23)) {
eval(parse(text =paste0("group",i,"dup<- subset (relatedness, Group=='",i,"' & PairGroup=='",i,"')")))
eval(parse(text =paste0("group",i,"<- remov_pwisedup(group",i,"dup)")))
}

#Groupmean generating pdf
pdf("Fig_AB_groupmean.pdf", family="Helvetica",width =11, height = 7)
boxplot(group1$r,group2$r,group3$r,group4$r,group5$r,group6$r,group7$r,group8$r,group9$r,group11$r,group12$r,group13$r,group14$r,group15$r,group16$r,group17$r,group18$r,group19$r,group20$r,group21$r,group22$r,group23$r, col="lightblue", border="#0080ff", las=1,names=c("#1","#2","#3","#4","#5","#6","#7","#8","#9","#11","#12","#13","#14","#15","#16","#17","#18","#19","#20","#21","#22","#23"),cex.axis=0.9)
mean_AB <- c(mean(group1$r),mean(group2$r),mean(group3$r),mean(group4$r),mean(group5$r),mean(group6$r),mean(group7$r),mean(group8$r),mean(group9$r),mean(group11$r),mean(group12$r),mean(group13$r),mean(group14$r),mean(group15$r),mean(group16$r),mean(group17$r),mean(group18$r),mean(group19$r),mean(group20$r),mean(group21$r),mean(group22$r),mean(group23$r))
points( mean_AB, pch = 21, cex =1, bg = "black")
dev.off()

mean(mean_AB)
var(mean_AB)

#mean relatedness significance test of correlated factors \
#colony size
Colony <- as.integer(c(1:9,11:23))
related <- data.frame(Colony, mean_AB,stringsAsFactors = F )
related

colonysize <- read.csv("Colony sizes.csv")
colonysize
data <- merge(colonysize,related,by="Colony")

testResult <- cor.test(data$number.of.aphids, data$mean_AB)
testResult #t = 1.6806, df = 20, p-value = 0.1084

testResult2 <- cor.test(data$frequency, data$mean_AB)
testResult2 #t = 0.53878, df = 20, p-value = 0.596


#pair-background
data <- read.csv("data.csv")
data$r <- as.numeric(data$relatedness)
data$num <- as.numeric(data$num)
summary(data)
hbpairs <- subset(data, data$group==data$group_pair & data$headbutt=="Y" & data$headbutt_pair=="Y" & data$num==data$num_pair & data$outcome=="W")
datar <- hbpairs[,c('ID','group','ID_pair','r')]
sum(datar$r==1)
#mean and SD
mean(datar$r)
var(datar$r)
#clonal pairs
sum(datar$r==1)
#write.csv(datar, file="datar.csv")

#bootstrap test
tt<-numeric(0)
for(i in 1:10000)　{
d <- numeric()
for (k in c(1:4,7:9,11:15,17:23))
{
eval(parse(text =paste0("x <- subset(datar,group=='",k,"')")))
eval(parse(text =paste0("y <- filter(group",k,",str_detect(ID, 'N') & str_detect(PairID, 'N'))")))
#print(y)
z <- sample(y$r,nrow(x),replace = TRUE)
#print(z)
d <- append(d,z)
}
tt[i] <- mean(d)
}
mean(tt)
mean(datar$r)
print(sum(tt > mean(datar$r))/10000 *2 ) #P=0.59, two-tailed

pdf("boot_hist.pdf", family="Helvetica",width =8, height = 7)
hist(tt, main="",xlim=c(0.5,1),xaxs="i", yaxs="i",ylim=c(0,2000),xlab="Bootstrapped value of relatedness", las=1, cex.axis=1.5, cex.lab=1.5) # expected distribution
abline(v=mean(datar$r), col="red",lty=2, lwd=2)
dev.off()


##permutation test - attackerwin vs defenderwin
hbpairs_field <- read.csv("hbpairs_field.csv")

attackerwin <- subset(hbpairs_field, hbpairs_field$won.by.attacker =="1" & relatedness!='NA')
defenderwin <- subset(hbpairs_field, hbpairs_field$won.by.attacker =="0" & relatedness!='NA')
summary(attackerwin)
summary(defenderwin)

d <- rbind(attackerwin,defenderwin)

mean(d$relatedness)
mean(attackerwin$relatedness)
sd(attackerwin$relatedness)
mean(defenderwin$relatedness)
sd(defenderwin$relatedness)

diff <- mean(attackerwin$relatedness) - mean(defenderwin$relatedness) #0.0545
set.seed(777)
tt<-numeric(0)
for(i in 1:10000)　{
picked = sample(seq_len(nrow(d)),size = nrow(attackerwin))
at <- d[picked,]
de <- d[-picked,]
mean(at$relatedness)
mean(de$relatedness)
tt[i] <- mean(at$relatedness)-mean(de$relatedness)
}
mean(tt)
print((sum(diff < tt)+sum(-diff >tt))/10000)  # p = 0.5009
hist(tt, main="",xlim=c(-0.4,0.4),xaxs="i", yaxs="i",ylim=c(0,3000),xlab="Permutation distribution", las=1, cex.axis=1.5, cex.lab=1.5) # expected distiribution
abline(v=diff, col="red",lty=2, lwd=2)
abline(v=-diff, col="red",lty=2, lwd=2)
```
